# TMS-induced inhibition of the left premotor cortex modulates illusory social perception

**DOI:** 10.1101/2023.01.30.526257

**Authors:** Charline Peylo, Elisabeth F. Sterner, Yifan Zeng, the EMPRA students, Elisabeth V. C. Friedrich

**Affiliations:** Department of Psychology /Research Unit Biological Psychology; Ludwig-Maximilians-Universität München; Munich, Bavaria, 80802; Germany; Department of Diagnostic and Interventional Neuroradiology /School of Medicine; Technical University of Munich; Munich, Bavaria, 81675; Germany

**Keywords:** Continuous Theta Burst Stimulation (cTBS), action prediction, interpersonal predictive coding, point-light agents, dyadic social interaction, communicative gestures

## Abstract

Communicative actions from one person are used to predict another person’s response. However, in some cases, these predictions can outweigh the processing of sensory information and lead to illusory social perception such as seeing two people interact, although only one is present (i.e., seeing a Bayesian ghost).

We applied either inhibitory brain stimulation over the left premotor cortex (i.e., real TMS) or sham TMS. Then, participants indicated the presence or absence of a masked agent that followed a communicative or individual gesture of another agent.

As expected, participants had more false alarms (i.e., Bayesian ghosts) in the communicative than individual condition in the sham TMS session and this difference between conditions vanished after real TMS. In contrast to our hypothesis, the number of false alarms increased (rather than decreased) after real TMS.

These pre-registered findings confirm the significance of the premotor cortex for social action predictions and illusory social perception.

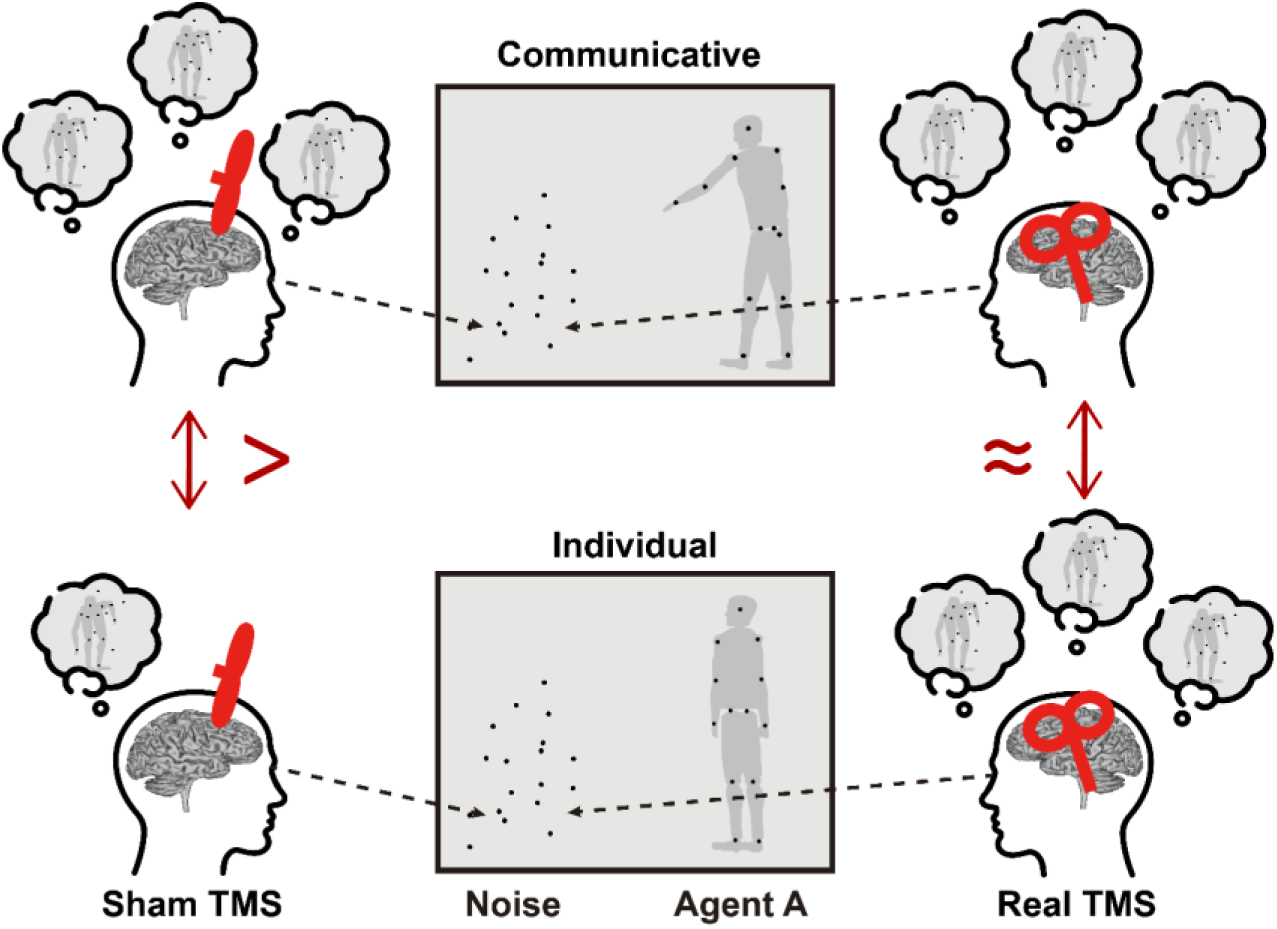

**Highlights:**

- Social predictions can outweigh sensory information and lead to illusory perception
- Premotor cortex is linked to the illusory social perception of a Bayesian ghost
- TMS over premotor cortex modulates how social predictions influence our perception

## Introduction

Our prior expectations affect our perception of others’ actions in social interactions. For example, you see your friend sticking out his hand to a woman as if he would like to shake hands and introduce himself. Due to your prior experience, you expect the woman to respond to the handshake. The use of prior experience to anticipate the interaction partner’s response action is called interpersonal predictive coding (Manera et al., 2011a, b; von der Lühe et al., 2016). Usually, interpersonal predictive coding helps us to handle complex social situations, but sometimes it can also lead to erroneous perception. Imagine in our example above that you are in a dark and smoky club when you see your friend trying to introduce himself to the woman. When you see the woman sticking out her hand too, you might have an illusion of them shaking hands when in fact the woman was just grabbing her drink and did not respond to your friend at all. This example should illustrate that our top-down expectations can be so strong that they overwrite our sensory information in ambiguous situations and lead us to perceive something or even someone who is not present. This phenomenon was called “seeing a Bayesian ghost” in previous work (Friedrich et al., 2022a; Manera et al., 2011b; von der Lühe et al., 2016). We use this term to describe an illusory perception of a person responding to a communicative gesture from another person due to prior expectations. The goal of this study was to investigate if there is a causal relationship between premotor activation and the occurrence of the Bayesian ghost.

We used an experimental design developed by Manera and colleagues (2011b) including two point-light agents (Figure 1; Friedrich et al., 2022a, b; Manera et al., 2011b; Zillekens et al., 2019). The agents are humans represented by moving lights that are attached to their major joints. Point-light displays are perceived as biological motion and can be used to investigate social interactions (Neri et al., 1998; Pavlova, 2012). In the current experimental design, agent A was well-recognizable and performed either a communicative or individual gesture. Agent B was blended into a cluster of noise dots and was either present (i.e., signal trials) or absent (i.e., noise trials). The participants’ task was to indicate whether they saw agent B in the cluster of noise dots or not. Previous studies using the same experimental design showed that communicative gestures, in contrast to individual actions, facilitated the perception of a second agent (Friedrich et al., 2022b; Manera et al., 2011b; Zillekens et al., 2019).

**Figure 1.**
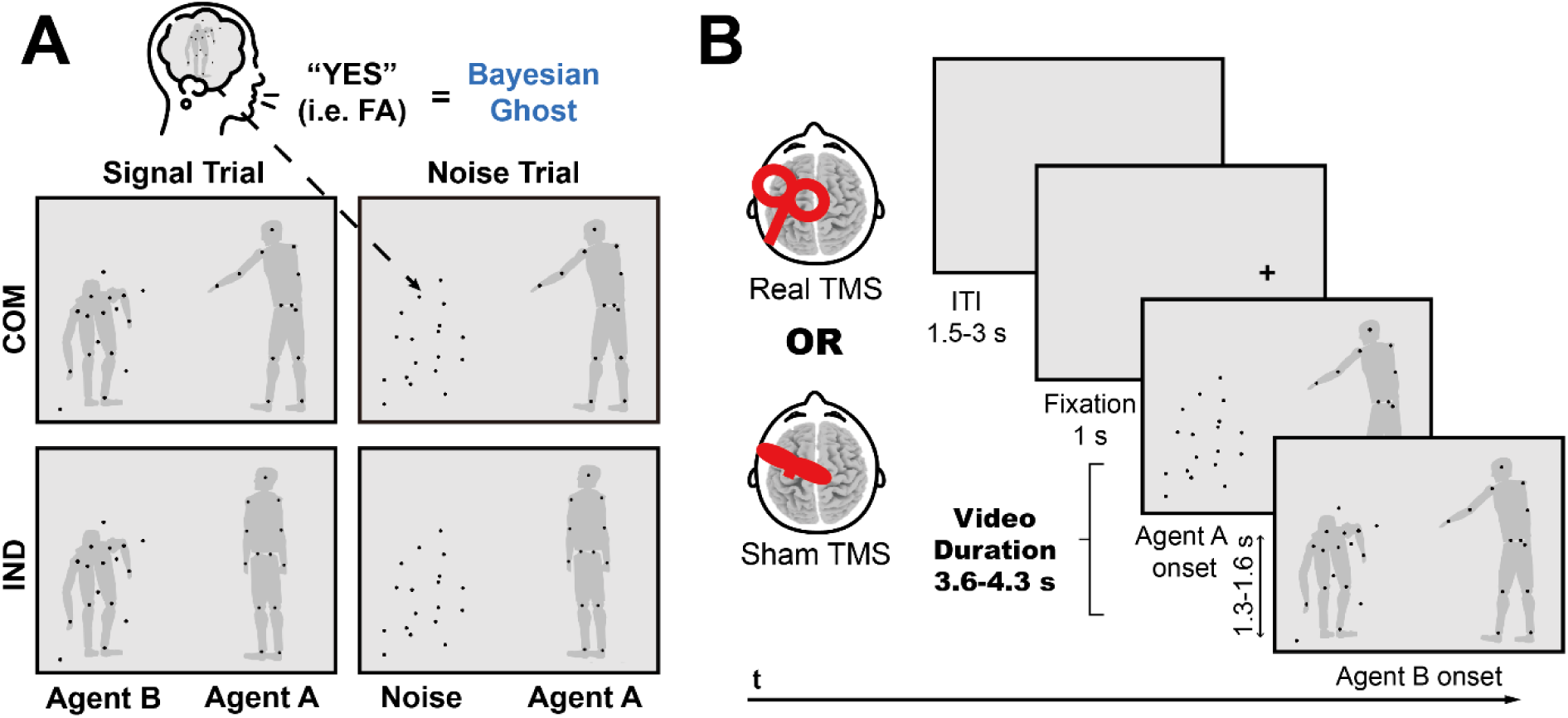
Experimental design (adapted from Friedrich et al., 2022a, b and Zillekens et al., 2019). **(A) Task conditions:** A well-recognizable agent A performed either a communicative (COM) or an individual (IND) gesture. Agent B was blended into a cluster of randomly moving noise dots and performed a gesture in response of agent A in half of the trials (i.e., signal trials). In the other half of the trials, agent B was entirely replaced by the noise dots (i.e., noise trials). The task of the participants was to report the presence or absence of agent B. The Bayesian ghost was defined as a false alarm (i.e., FA) in the communicative noise trial. Please note that the gray silhouettes only serve illustration purposes and were not visible for the participants. **(B) Timing:** The main experiment was divided into two sessions on two separate days. In each session, participants received either offline real or sham TMS. Following the TMS application, participants performed the task. Each trial was initiated with a jittered intertrial interval (i.e., ITI) between 1.5 - 3 s, followed by a 1 s fixation cross indicating the subsequent location of agent A. Then, the video displaying agent A and the cluster of dots (with or without agent B) started. Although, agent B was present from the beginning of the video, its response action was delayed between 1.3 -1.6 s. Participants were required to make a response as long as the video was still ongoing. The video duration dependent on the specific action and was between 3.6 - 4.3 s long. Please note that the gray silhouettes again only serve illustration purposes and were not visible for the participants.

Based on this behavioral finding, Manera and colleagues (2011b) concluded that interpersonal predictive coding in a social context can be so strong that it can lead to an illusion of a second agent, i.e., to seeing a “Bayesian ghost”. Von der Lühe et al. (2016) picked up on the term Bayesian ghost and showed that individuals on the autism spectrum did not exploit the biological motion signals in order to predict responses based on communicative intentions, although being able to explicitly recognize communicative intentions. These behavioral findings suggested a deficit in interpersonal predictive coding in Autism Spectrum Disorder. They are in line with the idea that individuals on the autism spectrum more strongly rely on sensory information and are less guided by predictions (Haker et al., 2016; Pellicano & Burr, 2012; Tarasi et al., 2022) and thus would see fewer Bayesian ghosts than neuro-typical controls.

To investigate the Bayesian ghost neuroscientifically, Friedrich and colleagues (2022a) asked neuro-typical participants to perform the above-described experimental task while recording their brain activity via electroencephalography (EEG). They defined the Bayesian ghost as a false alarm response in the communicative noise trial. These are trials, in which agent A performed a communicative gesture and the participants indicated to see agent B although it was not present in the cluster of noise dots. Their behavioral results showed that participants made more false alarms (i.e., participants indicated that agent B was present although it was absent) in the communicative than in the individual condition and the opposite was true for correct rejections (i.e., participants indicated correctly that agent B was absent). On the neural level, false alarms were compared to correct rejections within the communicative noise trials. The EEG results demonstrated that activation of the left premotor cortex during the observation of agent A’s communicative gesture predicted the occurrence of the Bayesian ghost in the later time segment. Importantly, the premotor activation was specific for this early time segment of the task, in which agent A was observed and the predictions for agent B’s response were formed. During the actual appearance of the Bayesian ghost (i.e., when participants watched the cluster of noise dots and indicated the presence or absence of agent B), more posterior regions in the sensorimotor cortex and adjacent parietal regions were activated.

These findings were in line with previous studies showing that the motor system is associated with the anticipation and mental simulation of future predictable actions (Cisek & Kalaska, 2004; Kilner et al., 2004; Krol et al., 2020; Maranesi et al., 2014; Umiltà et al., 2001; Urgesi et al., 2010). Moreover, these findings emphasized the importance of the left premotor cortex in order to generate strong top-down predictions that outweighed the sensory input in the later time segment and thus led to seeing the second agent although it wasn’t there, i.e., seeing the Bayesian ghost. The left premotor cortex was previously associated with the detection of biological motion (van Kemenade et al., 2012), action understanding (Michael et al., 2014) and action prediction (Brich et al., 2018; Stadler et al., 2012) in studies using non-invasive brain stimulation. The EEG results of Friedrich et al. (2022a) suggested that the left premotor cortex is associated with action prediction in a social context. Another brain region typically activated during the observation of social interactions (compared to individual actions), is the superior temporal sulcus (Caspers et al., 2010; Isik et al., 2017; Pavlova, 2012). However, as the Bayesian ghost is not defined by the difference between the communicative and individual conditions, we did not expect activation in the superior temporal sulcus to be crucial for our task. The Bayesian ghost occurs when a communicative gesture of one agent leads us to perceive the second agent, although it is not present (i.e., false alarm response in a communicative noise trial). Thus, we assumed that the Bayesian ghost is associated with action prediction in a social context, which was previously related to the left premotor cortex (Brich et al., 2018; Friedrich et al., 2022a; Stadler et al., 2012).

To test whether the premotor cortex is causally linked to the occurrence of the Bayesian ghost, transcranial magnetic stimulation (i.e., TMS) can be used (Walsh & Cowey, 2000). Depending on the specific TMS protocol, the effects of TMS can be excitatory or inhibitory, and short or long-lasting (di Lazzaro et al., 2011; Hallett, 2007; Huang et al., 2005).

Continuous Theta Burst Stimulation (cTBS) was shown to inhibit the motor cortex for up to an hour (Huang et al., 2005). Thus, we applied offline cTBS to inhibit the left premotor cortex right before participants started the experimental task. In a different session, we applied sham TMS to the left premotor cortex as control condition in a within-subject design with randomized session order (Figure 1).

If the premotor cortex is causally linked to the occurrence of the Bayesian ghost, we expect that inhibition of the left premotor cortex with cTBS will disrupt the processes involved in the formation of the Bayesian ghost. This might lead to a reduction in the number of false alarms in the communicative condition. Thus, in the real TMS session, there should be no difference between the number of false alarms in the communicative condition (because predictions are disrupted) and the individual condition anymore (because no predictions are possible in the first place). In contrast, in the sham TMS session, we expect to replicate the results from the study of Friedrich and colleagues (2022a): Participants should make more false alarms in the communicative condition (because predictions are made about agent B’s response action) than in the individual condition (because no predictions are possible).

## Results

### TMS effects are largely unaffected by participants’ beliefs about the type of stimulation

In order to make sure that participants were blind to the effects of the offline TMS application, we asked them at the end of the study how they had experienced stimulation in each session separately. For the real TMS session, seven participants thought an excitatory TMS has been applied, three thought it had been inhibitory and nine had taken it as sham. For the sham TMS session, three and four participants guessed excitatory and inhibitory, respectively, whereas twelve participants correctly took it as sham. No one guessed the TMS effects for both sessions correctly. Thus, it is very unlikely that participants’ beliefs biased our TMS results.

### Global performance is comparable across TMS sessions

Overall, participants performed equally well in both TMS sessions (sham: *M*=72.52%, *SD*=9.23%; real: *M*=71.51%, *SD*=9.44%; *t*(18)=0.42, *p*=.682, *d*=0.096; *BF_01_*=3.90) and required an equal amount of noise dots to achieve the desired average accuracy of approximately 70% (sham: *M*=18.00, *Mdn*=14.00, *SD*=14.08; real: *M*=22.32, *Mdn*=14.00, *SD*=28.00; *z*=-0.04, *p*=.983, *r_rb_*=-.012 (corresponding to a Cohen’s *d* of *d*=-0.024 according to Rosenthal, 1994, p. 239); *BF_01_*=4.20).

### TMS-induced inhibition of the left premotor cortex modulates the number of false alarms (i.e., Bayesian ghost occurrence)

The pre-registered 2x2x2 repeated-measures ANOVA for the number of responses with factors ‘TMS’ (real vs. sham), ‘condition’ (communicative vs. individual) and ‘response’ (false alarm vs. correct rejection) yielded a highly significant but trivial main effect of factor ‘response’ (*F*(1,18)=17.96, *p*<.001, η^2^_p_=.499; *BF_incl_*=28.11): As predicted based on the targeted 70% average accuracy, participants made overall more correct rejections (*M*=41.72≈65.63% of valid noise trials averaged across TMS sessions and task conditions, *SD*=13.13) than false alarms (*M*=20.71≈34.37% of valid noise trials averaged across TMS sessions and task conditions, *SD*=10.08). In addition to this main effect, we observed a significant interaction between factors ‘condition’ and ‘response’ (*F*(1,18)=7.75, *p*=.012, η^2^_p_=.301; *BF_incl_*=5.02), indicating a larger numerical advantage of correct rejections over false alarms in the individual condition (Δ*M*=23.63 averaged across TMS sessions, Δ*SD*=22.73) than in the communicative condition (Δ*M*=18.40 averaged across TMS sessions, Δ*SD*=21.24). The predicted three-way interaction between factors ‘TMS’, ‘condition’ and ‘response’ did not reach statistical significance (*F*(1,18)=2.17, *p*=.158, η^2^_p_=.108; *BF_incl_*=2.52), nor did any of the other effects (all *p*≥.124; all *BF_incl_*<0.99). Descriptive statistics underlying the absent three-way interaction however suggested that the difference between the number of false alarms in the communicative and the individual condition might have been reduced following real compared to sham TMS (sham: Δ*M*=3.90, Δ*SD*=6.84; real: Δ*M*=0.63, Δ*SD*=4.83), as originally predicted. And so was the condition difference between the number of correct rejections albeit to a smaller degree (sham: Δ*M*=-4.05, Δ*SD*=6.98; real: Δ*M*=-1.90, Δ*SD*=4.90). The results are summarized in Figure 2.

**Figure 2.**
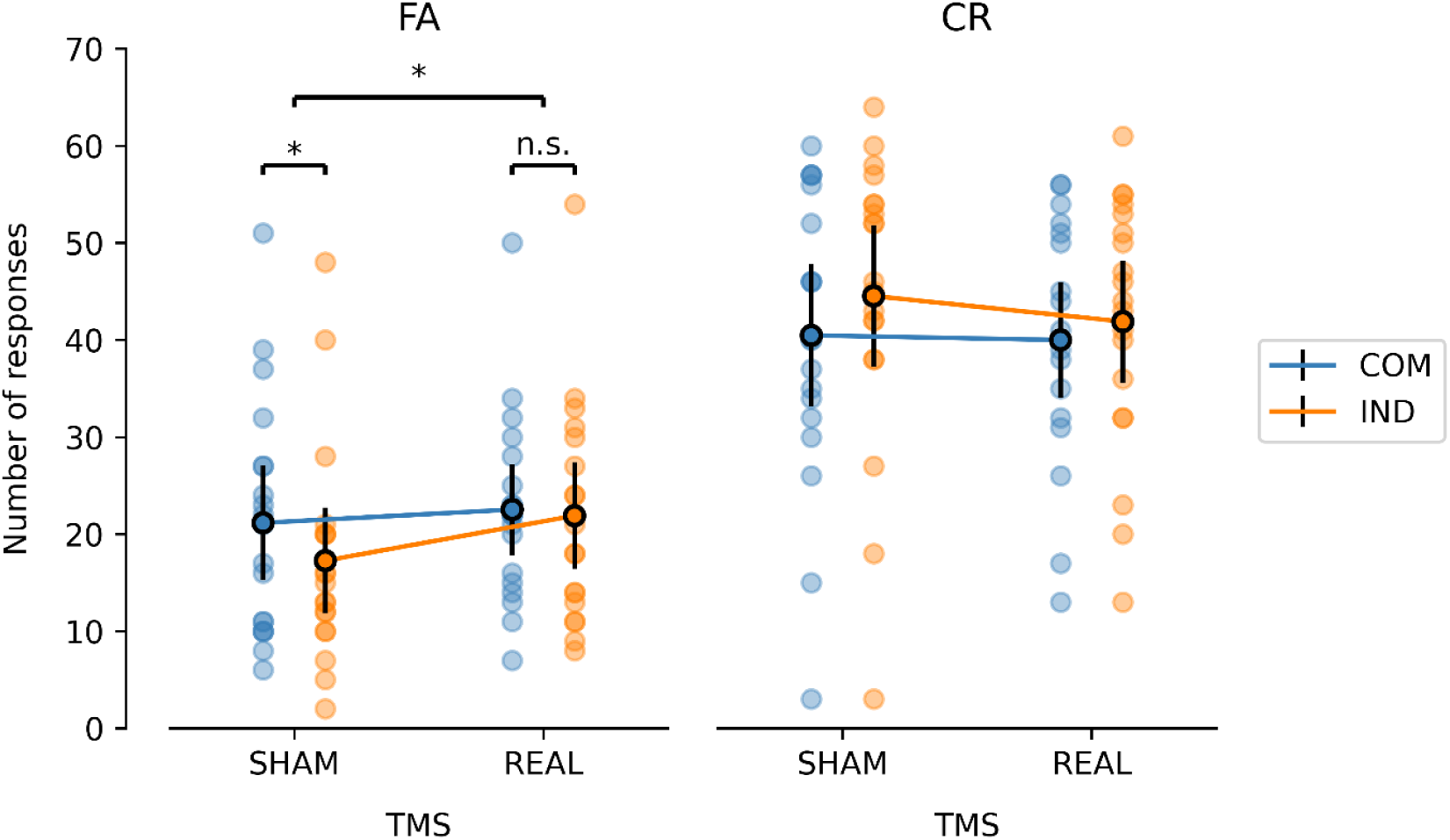
Real TMS eliminates the condition difference (COM-IND) for false alarms seen in sham TMS. Number of false alarms (FA; left column) and correct rejections (CR; right column) as a function of condition (communicative: COM vs. individual: IND) and TMS session (sham vs. real). Translucent dots represent the sampling distribution and opaque dots with error bars represent the sample mean and its 95% confidence interval. Asterisks highlight statistical significance at p < .05, whereas non-significant differences with p ≥ .05 are abbreviated with n.s.

To further explore our hypothesis that inhibition of the left premotor cortex through real TMS reduces the difference between the number of false alarms in the communicative and the individual condition, we contrasted the number of false alarms in the communicative and the individual condition separately for sham and real TMS. Additionally, we made a comparison of this condition difference (communicative - individual) between the two types of TMS. For these comparisons, we performed three additional Wilcoxon signed-rank tests and their Bayesian equivalents. In line with our hypotheses and previous results, in the sham TMS session, we observed significantly more false alarms in the communicative condition (*M*=21.16≈36.18% of valid noise trials, *Mdn*=21.00, *SD*=12.21) than in the individual condition (*M*=17.26≈29.82% of valid noise trials, *Mdn*=15.00, *SD*=11.32; *z*=2.20, *p_FDR_*=.045, *r_rb_*=.591 (corresponding to a Cohen’s *d* of *d*=1.465); *BF_10_*=9.72). Importantly, this condition difference was absent after applying real TMS (communicative: *M*=22.53≈36.66% of valid noise trials, *Mdn*=22.00, *SD*=9.76; individual: *M*=21.90≈34.76% of valid noise trials, *Mdn*=21.00, *SD*=11.34; *z*=0.54, *p_FDR_*=.600, *r_rb_*=.142 (corresponding to a Cohen’s *d* of *d*=0.287); *BF_01_*=3.43) and significantly reduced compared to sham TMS (sham: Δ*M*=3.90, Δ*Mdn*=3.00, Δ*SD*=6.84; real: Δ*M*=0.63, Δ*Mdn*=2.00, Δ*SD*=4.83; *z*=1.87, *p_FDR_*=.048, *r_rb_*=.516 (corresponding to a Cohen’s *d* of *d*=1.205); *BF_10_*=2.08). It should be noted, however, that this decreased condition difference following real compared to sham TMS was primarily driven by an increase in the number of false alarms in the individual condition (Δ*M*=4.63, Δ*Mdn*=4.00, Δ*SD*=7.82; *z*=2.29, *p_FDR_*=.046, *r_rb_*=.600 (corresponding to a Cohen’s *d* of *d*=1.500); *BF_10_*=6.67), rather than by the predicted decrease in the number of false alarms in the communicative condition (Δ*M*=1.37, Δ*Mdn*=-1.00, Δ*SD*=9.30; *z*=0.54, *p_FDR_*=.600, *r_rb_*=.142 (corresponding to a Cohen’s *d* of *d*=0.287); *BF_10_*=0.30).

### TMS exclusively affects illusory but not veridical social perception nor sensitivity

To check the specificity of the observed TMS effects for the Bayesian ghost occurrence as reflected by the number of false alarms (i.e., the detection of an agent although it is absent), we performed three control analyses. We contrasted sensitivity *d’* (i.e., the ability to distinguish signal from noise), criterion *c* (i.e., a potential bias towards one of the two response options ‘yes’ and ‘no’) and the number of hits (i.e., the correct detection of a present agent B) between the two TMS sessions (real vs. sham) and the two task conditions (communicative vs. individual) using three separate 2x2 repeated-measures ANOVAs.

In line with the descriptive statistics that were highly similar across all four conditions (sham- communicative: *M*=1.27, *SD*=0.68; sham-individual: *M*=1.43, *SD*=0.62; real-communicative: *M*=1.22, *SD*=0.64; real-individual: *M*=1.34, *SD*=0.68), the analysis of sensitivity *d’* yielded no significant effect of the factors TMS (*F*(1,18)=0.18, *p*=.676, η^2^_p_ =.010; *BF* =0.46), condition (*F*(1,18)=2.69, *p*=.119, η^2^_p_ =.130; *BF* =0.81) or their interaction (*F*(1,18)=0.14, *p*=.711, η^2^_p_ =.008; *BF_Incl_* =0.40).

The criterion *c* was also unaffected by the factor TMS (*F*(1,18)=1.10, *p*=.309, η^2^ =.057; *BF_Incl_*=0.65), but was (marginally) significantly affected by the factor condition (*F*(1,18)=3.75, *p*=.069, η^2^ =.172; *BF* =0.96) and its interaction with TMS (*F*(1,18)=4.97, *p*=.039, η^2^ =.216; *BF_Incl_*=2.19). Descriptive statistics indicated a stronger tendency towards ‘yes’ responses in the communicative compared to the individual condition following sham TMS (sham- communicative: *M*=-0.23, *SD*=0.54; sham-individual: *M*=-0.11, *SD*=0.59), which was absent after applying real TMS due to an increase of this ‘yes’ tendency in the individual condition (real-communicative: *M*=-0.24, *SD*=0.41; real-individual: *M*=-0.24, *SD*=0.44). It should be noted, however, that the normality assumption was violated for some variables and that the results of this analysis should thus be interpreted with caution.

Since the criterion is computed based on all ‘yes’ responses and could thus be driven either by the number of false alarms and/or by the number of hits, we also contrasted the latter between the two TMS sessions and the two task conditions. Similar to sensitivity *d’*, the number of hits was highly similar across all four conditions (sham-communicative: *M*=51.95≈76.42% of valid signal trials, *SD*=11.54; sham-individual: *M*=50.95≈74.40% of valid signal trials, *SD*=13.09; real-communicative: *M*=52.74≈77.13% of valid signal trials, *SD*=10.39; real-individual: *M*=53.37≈78.27% of valid signal trials, *SD*=9.82). Its analysis yielded no significant effect of the factors TMS (*F*(1,18)=0.33, *p*=.574, η^2^_p_=.018; *BF_Incl_*=0.58), condition (*F*(1,18)=0.03, *p*=.864, η^2^_p_=.002; *BF_Incl_*=0.35) or their interaction (*F*(1,18)=1.93, *p*=.182, η^2^_p_=.097; *BF_Incl_*=0.68).

## Discussion

### Bayesian ghost is linked to premotor cortex

This pre-registered study aimed to investigate if the occurrence of the Bayesian ghost is causally linked to premotor activation. In a within-subject design, we used real TMS (i.e., cTBS) versus sham TMS to inhibit the left premotor cortex offline before participants performed a point-light agent discrimination task. Although most of the participants distinguished the sham TMS from the real TMS application, it has to be noted that this question was only asked after completing both sessions, thereby facilitating a comparison between the two sessions. Only three out of 19 participants thought that the real TMS was inhibitory and none of the participants indicated the effect of both sessions correctly. Thus, we can assume that our experimental manipulation was successful. Moreover, these results suggest that any observed TMS effects should not be biased towards the predicted direction (i.e., reduced performance and/or Bayesian ghost perception) by participants’ beliefs about the TMS stimulation. Importantly, TMS was applied offline, i.e., there was no difference between the two sessions while participants were performing the experimental task.

In our pre-registration, we hypothesized that the difference between the communicative and individual conditions seen in the sham TMS session would disappear in the real TMS session (cf. ‘data analysis’ in https://osf.io/4fvns/?view_only=22773756c68c4d47b7ed27a25f3219ce). This was indeed the case in our data: In the sham TMS session, we could replicate the findings of Friedrich and colleagues (2022a): After seeing a communicative gesture of agent A, participants saw a Bayesian ghost (i.e., indicated by a false alarm in the communicative noise trial) more often than in the individual condition. In the real TMS session, the difference between the communicative and individual condition in the number of false alarms vanished. Additionally, we computed three control analyses in order to confirm that this effect was specific to the Bayesian ghost occurrence. Based on the response criterion, we found that the communicative actions facilitated the perception of a second agent compared to the individual condition in the sham TMS session (Friedrich et al., 2022b; Manera et al., 2011b; Zillekens et al., 2019), whereas this effect vanished after real TMS. However, neither our control analysis of sensitivity (targeting the overall ability to distinguish signal from noise) nor our control analysis of the number of hits (targeting the ability to correctly detect a present agent B, i.e., a correct ‘yes’ response) provided evidence for any such effect. The observed TMS effects on response criterion are therefore most likely a consequence of the change in the number of false alarms reported above and not a mere increase in ‘yes’ responses.

Together, these results suggest that TMS-induced inhibition of the left premotor cortex did not affect performance globally (e.g., by modulating the ability to distinguish signal from noise or to detect agent B if it was indeed there), but rather specifically targeted Bayesian ghost occurrence.

Thus, real TMS affected the processes underlying the illusory social perception of a second agent. This result is also in line with previous studies showing that the premotor cortex is involved in action predictions (Brich et al., 2018; Cattaneo & Parmigiani, 2021; Cisek & Kalaska, 2004; Kilner et al., 2004; Krol et al., 2020; Maranesi et al., 2014; Stadler et al., 2012; Umiltà et al., 2001; Urgesi et al., 2010). We can therefore extend the role of the premotor cortex for action prediction in a social context.

In contrast to our hypothesis, this elimination of differences between conditions was not due to a decrease in false alarms in the communicative condition but due to an increase in false alarms in the individual condition. We believe that real TMS over the premotor cortex led to a general increase in false alarms (e.g., van Kemenade et al., 2012) – which we did not consider in our pre-registration. If there was no influence of the real TMS premotor application specifically on the Bayesian ghost, there should still be the same difference between the communicative and individual conditions after real TMS that was shown in the sham session, only at a higher level. In contrast, this increase was seen in the individual condition, but was counteracted in the communicative condition, which still speaks for a specific link between the premotor cortex and the Bayesian ghost.

In the following chapter, we discuss different explanations for this result.

### Real TMS increased rather than decreased false alarms

First, we want to explain our assumption of real TMS introducing general noise, which became significant in the individual but counteracted in the communicative condition:

Assuming that the difference between the communicative and individual conditions lies in the additional effect arising from the predictive information inferred from agent A, TMS is unlikely to have only affected the individual but not the communicative condition. This is because the cognitive processes involved in the individual condition are a subset of those involved in the communicative condition. The present results could be explained by speculation that TMS influences prediction differently at different levels. At the low level, the premotor cortex generates predicted kinematics, including the precise timing of future movement sequences (Stadler et al., 2012). It might be utilized for both, communicative and individual, conditions in binding dots into a coherent biological movement. Real TMS did not necessarily abolish all the low-level prediction, but instead produced more prediction errors via an increased neural noise level (Miniussi et al., 2010; Ruzzoli et al., 2010; Schwarzkopf et al., 2011). It has been demonstrated that injecting low-level noise may raise the probability of detection by randomly pushing some of the weak signals above the threshold (Moss et al., 2004). In the noise trials of the present study, the movement of the random dots might be accidentally boosted by neural noise and erroneously interpreted as a body motion. In other words, the prediction was biased towards more false alarms. This is in line with a previous study where more false alarms of detecting point-light agents were reported after TMS over the premotor cortex (van Kemenade et al., 2012). Similarly, Stadler and colleagues (2012) as well as Brich and colleagues (2018) reported more prediction errors and less prediction accuracy, respectively, when inhibiting the left premotor cortex with TMS.

However, the false alarms remained unchanged in communicative condition after real TMS in the present study. This may relate to a higher semantic level of prediction specifically involved in the communicative condition. The premotor cortex was found to hold a goal-specific action repertoire (Rizzolatti & Fadiga, 1998) and exploit the corresponding action codes according to specific contexts (Schubotz et al., 2014). In this account, the communicative gesture may bias the activation of responsive action codes and generate a prediction of sequential action events as the template to extract more “meaningful” sequences (Hommel et al., 2001; Schubotz & Von Cramon, 2002) than it should. This leads to the discrepancy of false alarms between the communicative and individual condition, i.e., the Bayesian ghost. TMS may potentially disrupt this process and result in a decrease in false alarms in the communicative condition. However, as discussed above, this was compensated for by the increased false alarms via TMS’s effect on low-level prediction. Hence, no significant difference was found between real and sham TMS in the communicative condition.

Second, another possible explanation for the increase in the number of false alarms in the individual condition following real TMS might be a disturbance in the perception of agent A (Candidi et al., 2008; Caspers et al., 2010; Cross et al., 2009; Wheaton et al., 2004) rather than a disruption in the prediction of agent B. If real TMS decreased the participants’ ability to distinguish between the communicative and individual gestures (i.e., creating a situation of uncertainty) and biased them into mistaking individual for communicative ones, then that might explain why participants made more false alarms following real TMS (especially in the individual condition). In line with this reasoning, it has been shown that social judgments like emotion identification are adversely affected by uncertainty (Clarke et al., 2005) and that perceptual ambiguity can be associated with a bias towards social interpretations (Salge et al., 2020; Varrier & Finn, 2022). The opposite effect, however, can sometimes be observed as well (Vanrie et al., 2004). It should also be noted that participants were familiarized with the different gestures at the beginning of each session and that the set from which the stimuli in the present study originated has previously been shown to yield a high accuracy in communicative gesture identification and discrimination (Manera et al., 2010). Therefore, a TMS-induced disturbance in the prediction of agent B might provide a more plausible explanation for the increase in false alarms following TMS than a disrupted perception of agent A.

Third, another aspect to consider is that the premotor cortex is not the only region involved in action prediction and also not isolated within the brain but strongly interconnected with other areas. Also the parietal cortex was identified as core region for action predictions (Fontana et al., 2012). The premotor cortex and the parietal cortex are strongly interconnected and both are involved in action prediction (Balser et al., 2014; Ptak et al., 2017; Schubotz, 2007) by supporting predictive functions in the absence of visual input (Stadler et al., 2012). If the premotor cortex is inhibited, we could speculate that parietal areas are missing important input and thus produce erroneous output like illusory perception, independently of the condition. This speculation is supported by the results of Friedrich and colleagues (2022a) who suggested that the superior parietal region was involved in creating the illusion in absence of actual biological motion. Also in clinical studies, the superior parietal cortex was found to show impaired functional connectivity (Schilbach et al., 2016; Tarasi et al., 2021) and altered brain structures (Cachia et al., 2014; Delli Pizzi et al., 2014) in individuals with symptoms of hallucinations.

We cannot determine with certainty which of these explanations is the real cause of the increase in the number of false alarms after real TMS especially in the individual condition. However, it seems that TMS generally boosted the level of false alarms (as seen in the individual condition and reported in the literature, e.g. van Kemenade et al., 2012). As this increase in the number of false alarms was not significant in the communicative condition (i.e., the difference between the individual and communicative conditions disappeared after real TMS), we conclude that the inhibition of the left premotor cortex did impact the occurrence of the Bayesian ghost (Brich et al., 2018; Friedrich et al., 2022a; Stadler et al., 2012).

### Premotor cortex involvement in Bayesian ghosts most likely includes - but is not limited to - action prediction in a social context

In this study, the premotor cortex was inhibited offline with cTBS, which means that a virtual lesions was set before the participants started with the experimental task and this virtual lesion should have lasted for the whole session (Huang et al., 2005). The experimental task is rather complex and each trial consists of different parts, which can be divided into three time segments (Friedrich et al., 2022b): First, participants were watching agent A’s gesture and in case of a communicative gesture, they generated predictions for agent B’s response action. Second, they focused on the cluster of noise dots and tried to detect agent B among them. Third, they decided, whether they see agent B or not and indicated their choice with a button press. The inhibition of the premotor cortex could have different influences on this range of processes.

In the first time segment, an inhibited premotor cortex could have led to an impaired perception of agent A’s gesture (i.e., participants were not able any more to discriminate between individual and communicate gesture; also see above). This idea is supported by the study of Michael and colleagues (2014) who showed that participants were less accurate in recognizing pantomimed actions after inhibiting the respective area in the premotor cortex.

Thus, inhibiting the premotor cortex decreased performance in action understanding.

Another possibility is that participants could still recognize agent A’s gesture correctly but failed to generate or maintain (correct) predictions based on its communicative gesture. Premotor cortex activation was found to be especially involved in the initial phase of action prediction, i.e., to initiate the generation of predictions (Stadler et al., 2012; Urgesi et al., 2010). This is in line with the EEG study of Friedrich and colleagues (2022a) showing that premotor activation in the first time segment predicted the occurrence of the Bayesian ghost, which appeared in a later time segment. During the appearance of the Bayesian ghost, more posterior regions were activated. However, an inhibited premotor cortex could also influence the ability to observe biological motion in the cluster of noise dots in this later time segment (Fadiga et al., 2005).

As we used offline TMS, we cannot conclude which aspect(s) of the tasks were exactly targeted, but it might be more than one.

### Conclusions and relevance of the findings

Our findings show that the premotor cortex is linked to the illusory social perception of a Bayesian ghost, which is likely due to disrupted action predictions in a social context.

However, more studies are needed to control for possible and complex TMS effects such as the stimulation introducing unspecific noise or the effects spreading to whole brain networks.

Future studies could also apply TMS online to target the different neural dynamics involved in this complex task (Friedrich et al., 2022b). For example, one could administer TMS pulses during each trial to disrupt the neural processes either of the first time segment (i.e., viewing agent A’s communicative gestures and generating predictions about agent B’s action) or of the later time segment (i.e., viewing the cluster of dots and deciding whether agent B was present or not). This might allow conclusions to whether the premotor cortex is rather responsible for generating action predictions, observation of movements or both. Thus, we could determine more precisely the role of the premotor cortex in interpersonal predictive coding and the phenomenon of the Bayesian ghost.

Since illusory perception and impairments in social functioning are core symptoms of a variety of psychiatric disorders (Schilbach, 2016), our findings are also of clinical significance. Related to this, predictive coding has been proposed as a unifying framework to account for symptoms in Autism and Schizophrenia Spectrum Disorders (Tarasi et al., 2022). According to the framework of predictive coding, predictions (i.e., prior beliefs) are compared with incoming sensory information to make inferences about the external world. Critically, individuals with Autism or Schizophrenia Spectrum Disorders may rely on an imbalance in the weighting of prior beliefs and sensory evidence during this process (e.g., Adams et al., 2013, Pellicano & Burr, 2012; Sterzer et al., 2018; Van de Cruys et al., 2014). Additionally, the alterations in the disorders appear to be diametrically opposed, suggesting an autism-schizophrenia continuum (Tarasi et al., 2022): While positive symptoms in schizophrenia are driven by an overweighting of high-level prior beliefs (Corlett et al., 2019), symptoms in autism are linked to an overweighting of the sensory evidence (Haker et al., 2016). In relation to our paradigm, this is supported by research showing that high-functioning autistic individuals do not implicitly use social context to form predictions in order to enhance the detection of agent B in a noisy environment (von der Lühe et al., 2016). Although the present results do not directly infer a relationship between clinical symptoms and premotor cortex dysfunction, they do provide new insights into identifying brain regions responsible for generating action predictions in a social context and producing illusory social perception which is crucial for a better understanding of mental disorders and, ultimately, for improving treatment options.

### Limitations of the study

First, the TMS application had some limitations.

A. We compared sham versus real TMS using a within-subject design. For sham TMS, the coil was tilted by 90° in an upright position. Although, similar clicking sounds were audible, the coil pressure and the electrical tap perception on the scalp were different between real and sham TMS. However, as we applied TMS offline, there was no difference during the task performance between sham and real TMS.

After the last session, we asked participants to indicate for each session separately whether they felt the session excitatory, inhibitory or neutral. This kind of question could have biased participants to think that the sessions had a specific impact. However, as we asked the participants only after the last session, they were not biased during the experiment.

Moreover, participants could not tell the intended direction of effects. Future studies could additionally use visual analogue scales to rate the participants’ levels of comfort/discomfort caused by TMS (e.g. as used in Conde et al., 2019).

(B) TMS effects can be complex and the causal chain can be influenced by many confounding factors (Bergmann & Hartwigsen, 2021), making it difficult to reveal a direct causal relationship between brain and behavior. Moreover, the effects of TMS can depend on the physiological state of the brain and predisposition of the participants (Siebner et al., 2004). While some studies found stable inhibitory effects of cTBS (Romero et al., 2022), also low reproducibility and opposite effects of theta burst stimulation were reported (Ozdemir et al., 2021). Future studies should additionally include an inhibitory stimulation of a control site (other than the premotor cortex) or an excitatory stimulation of the premotor cortex (e.g., with iTBS) in order to control for TMS-induced noise and to further support the claim of causality.
(C) Another limitation of our TMS design was that we did not use neuronavigation in order to enhance the spatial precision. Future studies should use individual brain scans and neuronavigation technology to precisely target the same premotor coordinates for each participant.
(D) As already mentioned in the discussion section, our experimental design targeted multiple cognitive processes from biological motion perception, over gesture discrimination to action prediction; all of which are critical at different points in time and potential explanations for our results. Since our offline application of TMS most likely induced a general rather than a time- and thus process-specific inhibition of the premotor cortex, the question of the mechanism underlying our findings remains open to discussion.

Second, it should be noted that only the exploratory tests but not the planned ANOVA yielded significant effects of TMS on task performance. A potential reason for this difference is that the follow-up tests focused only on the number of false alarms (in line with the definition of a Bayesian ghost as an illusory percept of a non-existent second agent; i.e., a false alarm), whereas the ANOVA also included the number of correct rejections. The rationale behind this was to match the methods between the current experiment and the preceding study by Friedrich and colleagues (2022b), allowing for a direct comparison between our results and the ones in support of premotor cortical involvement in the occurrence of a Bayesian ghost. However, since the number of correct rejections was neither central to our hypotheses, nor to the effect size estimation underlying our a-priori power analysis, this factor might have just added more variance to the model without providing additional explanatory power. This might have led to a smaller overall effect size than what we had anticipated based on the false alarm literature alone and hence to a lower statistical power; an issue that might have been potentiated by the type of power analysis that we performed (“GPower 3.0”), which is less conservative than some of the other methods (e.g., “Cohen (1988)”). Despite the reduced power in non-parametrical testing, the exploratory tests focusing on the number of false alarms as central element of our hypotheses provided evidence that the illusory social perception of a Bayesian ghost indeed depends on the premotor cortex.

## Acknowledgments

We would like to thank Imme Zillekens, Leonhard Schilbach, Valeria Manera, Cristina Becchio and Atesh Koul for providing the experimental task and advice during setup and analysis. Moreover, our thanks goes to Paul Sauseng for providing valuable advice throughout the study and to Yannik Hilla for the supportive discussions regarding Bayesian statistics. This study was funded by a DFG grant to E.V.C.F. (FR 3961/1-1).

## Authors contributions

CP: Conceptualization, Methodology, Software, Validation, Formal analysis, Investigation, Data curation, Resources, Writing – original draft, Writing – review & editing, Visualization, Supervision, Project administration.

EFS: Investigation, Data curation, Project administration.

YZ: Writing – original draft, Writing – review & editing, Visualization. EMPRA students: Conceptualization, Methodology, Investigation.

EVCF: Conceptualization, Methodology, Software, Validation, Formal analysis, Investigation, Data curation, Resources, Writing – original draft, Writing – review & editing, Visualization, Supervision, Project administration, Funding acquisition.

## STAR METHODS

### RESOURCE AVAILABILITY

#### Lead contact

Further information and requests should be directed to and will be fulfilled by the lead contact, Elisabeth V. C. Friedrich.

#### Materials availability

This study did not generate new unique reagents.

#### Data and code availability

The behavioral data and analysis have been deposited at the Open Science Framework (https://osf.io/) and are publicly available as of the date of publication. DOIs are listed in the key resources table.

Any additional information required to reanalyze the data reported in this paper is available from the lead contact upon request.

### EXPERIMENTAL MODEL AND STUDY PARTICIPANT DETAILS

An a-priori power analysis was performed with G*Power 3.1 (Faul et al., 2009) to determine the sample size for a repeated-measures ANOVA (within factors) with α = 0.05, power of 1 – β = 0.8, one group, two measurements, correlation of 0.5 among repeated measures and nonsphericity correction of 1 (default option as in G*Power 3.0). Previous TMS literature reported effect sizes of d = 0.9 - 1.2 (Basil et al., 2017; Lapenta et al., 2022; Michael et al., 2014; Stadler et al., 2012; van Kemenade et al., 2012). Yet we suggest such effect sizes may be overestimated due to rather small sample sizes tested in these studies. As a conservative estimation, an effect size of d = 0.7 (f = 0.35) was used to find differences between sham and real TMS, and resulted in a minimal sample size of 19 participants.We recruited 21 young, neuro-typical participants. All of them were right-handed, had normal or correct-to-normal vision, no diagnosis of neurological or psychiatric disorders, no history of medication intake and were not pregnant. In addition, all fulfilled the TMS safety criteria in line with Rossi and colleagues (2009). Two participants were excluded: For one participant, the real TMS session had to be abandoned due to a fire alarm; and the other one failed to perform significantly above chance level in the sham TMS session (same criteria as in Friedrich and colleagues (2022a)). The remaining 19 participants (5 male and 14 female) were on average 21.6 (SD = 3.7) years old and obtained a mean Autism Spectrum Quotient of 14.1 (SD = 5.0; Baron-Cohen et al., 2001), which is indicative of a control group without clinically significant levels of Autism Spectrum Disorder. The participants gave written informed consent prior to the study and received course credits as compensation. The study was conducted under the approval of the LMU Munich Faculty 11 ethics committee and in accordance with the Declaration of Helsinki. The participants had no information about the different stimulation protocols (i.e., whether the TMS application was excitatory, inhibitory or sham) during the experiment.

### METHOD DETAILS

#### Pre-registration

The experiment was pre-registered on Open Science Framework (https://osf.io/4fvns/?view_only=22773756c68c4d47b7ed27a25f3219ce). The hypotheses were defined in the paragraphs ‘hypothesis’ as well as ‘data analysis’.

#### Experimental Task

We used the same experimental task as the one in Friedrich and colleagues (2022a). It was programmed with Matlab R2016a (The MathWorks, Inc., Natick, Massachusetts, United States) and Psychophysics Toolbox (Version 3.0.14; Brainard & Vision, 1997; Kleiner et al., 2007; Pelli & Vision, 1997).

The visual stimuli were presented on a 1280 × 1024 monitor with a refresh rate of 60 Hz. The video included two point-light agents, which were each defined by 13 moving dots and were displayed on the two sides of the gray screen (Figure 1a). Agent A was well-recognizable and performed one out of six possible actions that either revealed an intention to communicate (communicative condition) or not (individual condition). Agent B performed actions responding to agent A’s communicative gestures.

In the communicative condition, the following three interactions between the two agents were included:

1. agent A: asking to squat down – agent B: squatting down;
2. agent A: asking to look at the ceiling – agent B: looking at the ceiling;
3. agent A: asking to sit down – agent B: sitting down.

In the individual condition the following actions were performed:

1. turning around – agent B: squatting down;
2. sneezing – agent B: looking at the ceiling;
3. agent A: drinking – agent B: sitting down.

Agent B was blended into a cluster of moving noise dots in half of the trials (i.e., signal trials). In the other half of the trials, it was entirely replaced by noise dots (i.e., noise trials) – whereas agent A’s actions were the same as in the signal trials. Thus, agent A’s action determined whether it was a communicative or individual condition and the presence or absence of agent B determined whether it was a signal or noise trial. The participants’ task was to indicate whether agent B was present or absent in the cluster of noise dots.

The light-point stimuli were chosen from the Communicative Interaction Database (Manera et al., 2010). Manera and colleagues asked human actors to perform communicative interactions together as well as non-communicative actions alone. Using motion capture and animation software, these real human (inter)-actions were transferred into point-light stimuli (for more details, please see Manera et al., 2010).

The point-light agents constitutes of 13 dots to define their bodies. In signal trials for agent B, however, only 6 dot positions were occupied by signal dots at a given time. The dots disappeared and reappeared at another random position after 200 ms. This happened asynchronous between dots. The same limited lifetime technique was used for the noise dots, which were identical with the signal dots. However, the noise dots appeared in an area sustaining an angle of about 8.6° x 14.3°, i.e., they were temporally and spatially scrambled. In noise trials, agent B was completely substituted by limited lifetime scrambled dots. The same number of noise dots were added as in the signal trials, so signal and noise trials were perceptually similar. For more details, please see Manera et al. (2011a) and Manera et al. (2011b).

Each trial began with a blank screen for an inter-trial interval of 1.5-3 s and was followed by a 1 s fixation cross indicating the subsequent position of agent A on the left or right side of the screen, which was counterbalanced. Then, agent A was presented together with the cluster of moving dots. Agent A’s action started immediately with the onset of the video. If agent B was present in the cluster of dots, its response action was delayed between 1267 - 1567 ms from agent A’s onset. Depending on the specific action, the videos lasted for 3600 - 4300 ms. Participants were asked to first passively observe agent A and then, to watch the cluster of noise dots. They should decide as fast and accurate as possible whether agent B was present or absent and indicate their choice by pressing the dedicated Yes or No key on a keyboard. The keys were counterbalanced between yes and no responses. Participants had to respond while the video with the point-light agent(s) was still ongoing. As we stimulated the left premotor cortex, we asked all participants to make their choice with the left hand.

#### TMS Apparatus

We delivered continuous Theta Burst Stimulation (cTBS) offline via a Mag&More PowerMag Research 100 TMS stimulator (Mag&More®) with a 7 cm figure-eight coil (Mag&More®). We first determined the active motor threshold over the left primary motor cortex. Participants maintained a voluntary flexion of their right wrist while actively stretching out their fingers. Single pulses were delivered to their left primary motor cortex with increasing intensity until a motor response was observable in exactly 3 out of 6 trials. The motor response was not measured with electromyography but determined by visually detecting muscle movement. This ‘observation of movement’ method was reported to be reliable (Varnava et al., 2011). For the actual cTBS, the coil was moved from the primary motor cortex 1.5 cm in the anterior direction and placed on the premotor cortex. We followed the cTBS protocol described by Huang and colleagues (2005), which is the standard for cTBS applications and safe (Rossi et al., 2009, 2021). The stimulation was set to an intensity of 80% of the individual active motor threshold.

The active motor threshold was on average 45% of the maximal stimulator output. We applied a 40 s continuous train, including bursts of three pulses at 50 Hz repeating every 200 ms (600 pulses in total). The 40-s stimulation was applied offline, i.e., before the experimental task was performed. The sham TMS session followed the same procedure (i.e., determining the motor threshold over the left primary motor cortex and then moving the coil to the left premotor cortex while applying the 600 pulses with the same stimulation parameters offline with an intensity of 80% of the individual active motor threshold). But the difference between the sham and real TMS session was that the coil was tilted by 90 degrees in an upright position during the cTBS application so that no effect on the brain was induced while retaining the clicking sound (see Figure 1b).

#### Procedure and Study Design

Each participant took part in two sessions on different days, which were 2-9 days apart from each other. In one session real TMS and in the other one sham TMS was applied in counterbalanced order between participants. Before the experiment, participants were asked to sign the informed consent, provide their age and gender, fill out the Autism Spectrum Quotient (Autism Research Center; Baron-Cohen et al., 2001) and the short form of the Edinburgh Handedness Inventory (Veale, 2013). In addition, participants were screened according to the TMS safety criteria in line with Rossi and colleagues (2009) and were asked about their vision, possible neurological or psychiatric disorders, medication intake or pregnancy.

At the beginning of both sessions, we determined the active motor threshold of each participant to define the stimulation intensity used for the real or sham TMS application. Similarly, we individualized the task difficulty in both sessions with a pre-test including 108 trials of an established procedure (Friedrich et al., 2022a, b; Manera et al., 2011b; Zillekens et al., 2019): Participants were asked to identify the presence or absence of agent B, which was blended into a cluster of noise of either 5, 20, or 40 dots. We fitted the response curve with a cumulative Gaussian function and obtained the number of dots corresponding to an average performance of 70%. This served as the individually adapted number of noise dots in the main experimental task. We used a minimum of five noise dots, even when the participants’ estimated number of dots was lower (Manera et al., 2011b, 2011a; Zillekens et al., 2019). As the pre-test was not identical with the main experiment, participants had the opportunity to get familiar with the experimental task during twelve example trials.

Then, the 40-s real or sham TMS – depending on the session - was applied offline right before participants performed the main experimental task (Figure 1B). We recorded 288 trials per participant in the main experiment (i.e., 72 trials per communicative/individual conditions x signal/noise trials). In total, each of the 6 different agent A – B stimulus pairings were shown 48 times. Specifically, the three communicative and the three individual A-B stimulus pairings for the signal and noise trials were each shown 12 times with agent A on the right and 12 times with agent B on the left side of the screen (i.e., 3 A - B stimulus pairings x 2 conditions x 2 signal/noise trials x 2 sides of agent A x 12 times = 288). The trials were organized in four 9-min blocks with breaks in between.

One session - including the instruction, determination of the motor and noise-thresholds, the TMS application and the experimental task - lasted about 100 minutes.

At the end of the last session of the experiment, participants were asked to fill out a questionnaire to indicate separately for the two sessions whether they experienced the TMS applications as excitatory, inhibitory or sham.

### QUANTIFICATION AND STATISTICAL ANALYSIS

First, we analyzed the questionnaire data. Then, we analyzed the responses of the participants from the experimental task. We excluded trials in which a response was given before agent B’s onset plus 200 ms or after the agents had already disappeared (i.e., invalid trials) and only considered those trials that were answered between agent B’s onset plus 200 ms and the disappearance of both agents (i.e., valid trials).

We compared the two TMS sessions in terms of performance (percent correct) and the number of noise dots needed to achieve the desired accuracy of 70%. For this purpose, we computed a two-sided dependent *t* test (for the normally distributed percent correct as suggested by a non-significant Shapiro-Wilk test, *p_SW_*>.05) and a two-tailed Wilcoxon signed-rank test (for the non-normally distributed number of noise dots; *p_SW_*<.05) together with their Bayesian equivalents (using a default Cauchy prior of 0.707 and a Markov Chain Monte Carlo algorithm with 1000 samples).

Afterwards, we performed the pre-registered 2x2x2 repeated-measures ANOVA (https://osf.io/4fvns/?view_only=22773756c68c4d47b7ed27a25f3219ce) for the number of responses with factors ‘TMS’ (real vs. sham), ‘condition’ (communicative vs. individual) and ‘response’ (false alarm vs. correct rejection). For the sake of completeness and comparability between this and all following analyses, which were performed using both frequentist and Bayesian statistics, we complemented the preregistered classical ANOVA with its Bayesian equivalent (quantifying the evidence for observing the reported data under models including the above factors or their interactions compared to matched models without such using a uniform prior of 0.5). To account for the non-normality (*p_SW_*<.05) and presence of outliers (as suggested by visual boxplot inspection) in some of the above variables (which could potentially bias the ANOVA’s results), and to further explore our hypothesis that inhibition of the left premotor cortex through real TMS reduces the difference between the communicative and the individual condition in the number of false alarms, we performed three additional Wilcoxon signed-rank tests (to account for outliers in some of the variables) and their Bayesian equivalents (using a default Cauchy prior of 0.707 and a Markov Chain Monte Carlo algorithm with 1000 samples). These follow-up tests contrasted the number of false alarms in the communicative and the individual condition separately for sham and real TMS, as well as the condition difference (communicative – individual) between the two types of TMS. Because we expected (a) significantly more false alarms in the communicative than in the individual condition when applying sham TMS, (b) no such difference after applying real TMS and hence (c) a significant reduction of this condition difference following real compared to sham TMS, the first and last test were one-tailed and the second test was two-tailed. FDR correction was used to account for multiple comparisons and the associated increased risk of false positives. Descriptive statistics underlying these analyses indicated that the decreased condition difference was primarily driven by an increase in the number of false alarms in the individual condition rather than by the predicted reduction in the communicative condition (see *Results*). We therefore performed two additional two-tailed Wilcoxon signed-rank tests (due to outliers in some of the variables) together with their Bayesian equivalents (using a default Cauchy prior of 0.707 and a Markov Chain Monte Carlo algorithm with 1000 samples) to contrast the number of false alarms between the two TMS sessions separately for both task conditions (using FDR correction for multiple comparisons) to quantify the statistical significance of this observation.

Lastly, to check if the predicted TMS effects were indeed specific to the number of false alarms (i.e., the detection of an agent although it is absent), we performed three control analyses, in which we contrasted sensitivity *d’* (i.e., the ability to distinguish signal from noise), criterion *c* (i.e., a potential bias towards one of the two possible response options) and the number of hits (i.e., the correct identification of a present agent B) between the two TMS sessions (real vs. sham) and the two task conditions (communicative vs. individual) using three separate 2x2 repeated-measures ANOVAs and their Bayesian equivalents (quantifying the evidence for observing the reported data under models including the above factors or their interactions compared to matched models without such using a uniform prior of 0.5).

All statistical tests were performed using JASP 0.16.4 (JASP Team, 2022) and the corresponding data was plotted in the Spyder 4.1.5 environment (Spyder Developer Team, 2020) for Python 3.7.3 (Van Rossum & Drake, 2009) using custom-written scripts and the open-source packages Matplotlib 3.3.2 (Hunter, 2007) and NumPy 1.19.2 (Harris et al., 2020). Test results were considered statistically significant if the probability of the results being random (i.e., showing no differences between means) was less than 5% (i.e., if *p* or in the case of FDR correction *p_FDR_* were <.05). The descriptive and statistical details indicated by the mean (*M*), median (*Mdn*), standard deviation (*SD*), the number of participants (*N*), the effect sizes Cohen’s *d,* partial Eta squared (η^2^_p_) and the rank-biserial correlation coefficient (*r_rb_*), the *z* value (reflecting the standardized *W* parameter from the Wilcoxon signed-rank test) and the *t* and *F* values with their degrees of freedom (*t*(X) and *F*(X,X), respectively), as well as the H0-over-H1 (*BF_01_*), H1-over-H0 (*BF_10_*) and inclusion Bayes factors (*BF_incl_*) can be found in the *Results* section. In line with Lee and Wagenmakers (2014), Bayes factors from 1 to 3, 3 to 10 and 10 to 30 were considered anecdotal, moderate and strong evidence for the null hypothesis (*BF_01_*), the alternative hypothesis (*BF_10_*) and for the inclusion of a certain model factor (*BF_incl_*), respectively. Conversely, Bayes factors from 0.33 to 1, 0.1 to 0.33 and 0.033 to 0.1 were interpreted as anecdotal, moderate and strong evidence for the alternative hypothesis (*BF_01_*), the null hypothesis (*BF_10_*) and for the exclusion of a certain model factor (*BF_incl_*), respectively.

## References

Adams, R. A., Stephan, K. E., Brown, H. R., Frith, C. D., & Friston, K. J. (2013). The computational anatomy of psychosis. Frontiers in Psychiatry, 4, 47. https://doi.org/10.3389/FPSYT.2013.00047/BIBTEX

Balser, N., Lorey, B., Pilgramm, S., Stark, R., Bischoff, M., Zentgraf, K., Williams, A. M., & Munzert, J. (2014). Prediction of human actions: expertise and task-related effects on neural activation of the action observation network. Human Brain Mapping, 35(8), 4016– 4034. https://doi.org/10.1002/HBM.22455

Baron-Cohen, S., Wheelwright, S., Skinner, R., Martin, J., & Clubley, E. (2001). The Autism- Spectrum Quotient (AQ): Evidence from Asperger Syndrome/High-Functioning Autism, Males and Females, Scientists and Mathematicians. In Journal of Autism and Developmental Disorders (Vol. 31, Issue 1).

Basil, R. A., Westwater, M. L., Wiener, M., & Thompson, J. C. (2017). A Causal Role of the Right Superior Temporal Sulcus in Emotion Recognition From Biological Motion. Open Mind, 2(1), 26–36. https://doi.org/10.1162/OPMI_A_00015

Bergmann, T. O., & Hartwigsen, G. (2021). Inferring Causality from Noninvasive Brain Stimulation in Cognitive Neuroscience. Journal of Cognitive Neuroscience, 33(2), 195–225. https://doi.org/10.1162/JOCN_A_01591

Brainard, D. H., & Vision, S. (1997). The psychophysics toolbox. Spatial Vision, 10(4), 433– 436.

Brich, L. F. M., Bächle, C., Hermsdörfer, J., & Stadler, W. (2018). Real-time prediction of observed action requires integrity of the dorsal premotor cortex: Evidence from repetitive transcranial magnetic stimulation. Frontiers in Human Neuroscience, 12, 101. https://doi.org/10.3389/FNHUM.2018.00101/BIBTEX

Cachia, A., Amad, A., Brunelin, J., Krebs, M. O., Plaze, M., Thomas, P., & Jardri, R. (2014). Deviations in cortex sulcation associated with visual hallucinations in schizophrenia. Molecular Psychiatry 2014 20:9, 20(9), 1101–1107. https://doi.org/10.1038/mp.2014.140

Candidi, M., Urgesi, C., Ionta, S., & Aglioti, S. M. (2008). Virtual lesion of ventral premotor cortex impairs visual perception of biomechanically possible but not impossible actions. Https://Doi.Org/10.1080/17470910701676269, 3(3–4), 388–400. https://doi.org/10.1080/17470910701676269

Caspers, S., Zilles, K., Laird, A. R., & Eickhoff, S. B. (2010). ALE meta-analysis of action observation and imitation in the human brain. NeuroImage, 50(3), 1148–1167. https://doi.org/10.1016/J.NEUROIMAGE.2009.12.112

Cattaneo, L., & Parmigiani, S. (2021). Stimulation of Different Sectors of the Human Dorsal Premotor Cortex Induces a Shift from Reactive to Predictive Action Strategies and Changes in Motor Inhibition: A Dense Transcranial Magnetic Stimulation (TMS) Mapping Study. Brain Sciences 2021, Vol. 11, Page 534, 11(5), 534. https://doi.org/10.3390/BRAINSCI11050534

Cisek, P., & Kalaska, J. F. (2004). Neural correlates of mental rehearsal in dorsal premotor cortex. Nature, 431(7011), 993–996. https://doi.org/10.1038/NATURE03005

Clarke, T. J., Bradshaw, M. F., Field, D. T., Hampson, S. E., & Rose, D. (2005). The perception of emotion from body movement in point-light displays of interpersonal dialogue. Perception, 34(10), 1171–1180. https://doi.org/10.1068/p5203

Conde, V., Tomasevic, L., Akopian, I., Stanek, K., Saturnino, G. B., Thielscher, A., Bergmann, T. O., & Siebner, H. R. (2019). The non-transcranial TMS-evoked potential is an inherent source of ambiguity in TMS-EEG studies. NeuroImage, 185, 300–312. https://doi.org/10.1016/J.NEUROIMAGE.2018.10.052

Corlett, P. R., Horga, G., Fletcher, P. C., Alderson-Day, B., Schmack, K., & Powers, A. R. (2019). Hallucinations and Strong Priors. In Trends in Cognitive Sciences (Vol. 23, Issue 2). https://doi.org/10.1016/j.tics.2018.12.001

Cross, E. S., Kraemer, D. J. M., Hamilton, A. F. de C., Kelley, W. M., & Grafton, S. T. (2009). Sensitivity of the Action Observation Network to Physical and Observational Learning. Cerebral Cortex, 19(2), 315–326. https://doi.org/10.1093/CERCOR/BHN083

Delli Pizzi, S., Franciotti, R., Tartaro, A., Caulo, M., Thomas, A., Onofrj, M., & Bonanni, L. (2014). Structural Alteration of the Dorsal Visual Network in DLB Patients with Visual Hallucinations: A Cortical Thickness MRI Study. PLOS ONE, 9(1), e86624. https://doi.org/10.1371/JOURNAL.PONE.0086624

di Lazzaro, V., Dileone, M., Pilato, F., Capone, F., Musumeci, G., Ranieri, F., Ricci, V., Bria, P., Iorio, R. Di, de Waure, C., Pasqualetti, P., & Profice, P. (2011). Modulation of motor cortex neuronal networks by rTMS: Comparison of local and remote effects of six different protocols of stimulation. Journal of Neurophysiology, 105(5), 2150–2156. https://doi.org/10.1152/JN.00781.2010/ASSET/IMAGES/LARGE/Z9K0051107170001.J PEG

Fadiga, L., Craighero, L., & Olivier, E. (2005). Human motor cortex excitability during the perception of others’ action. Current Opinion in Neurobiology, 15(2), 213–218. https://doi.org/10.1016/J.CONB.2005.03.013

Faul, F., Erdfelder, E., Buchner, A., & Lang, A. G. (2009). Statistical power analyses using G*Power 3.1: Tests for correlation and regression analyses. Behavior Research Methods, 41(4), 1149–1160. https://doi.org/10.3758/BRM.41.4.1149/METRICS

Fontana, A. P., Kilner, J. M., Rodrigues, E. C., Joffily, M., Nighoghossian, N., Vargas, C. D., & Sirigu, A. (2012). Role of the parietal cortex in predicting incoming actions. NeuroImage, 59(1), 556–564. https://doi.org/10.1016/J.NEUROIMAGE.2011.07.046

Friedrich, E. V. C., Zillekens, I. C., Biel, A. L., O’Leary, D., Seegenschmiedt, E. V., Singer, J., Schilbach, L., & Sauseng, P. (2022a). Seeing a Bayesian ghost: Sensorimotor activation leads to an illusory social perception. IScience, 25(4), 104068. https://doi.org/10.1016/j.isci.2022.104068

Friedrich, E. V. C., Zillekens, I. C., Biel, A. L., O’Leary, D., Singer, J., Seegenschmiedt, E. V., Sauseng, P., & Schilbach, L. (2022b). Spatio-temporal dynamics of oscillatory brain activity during the observation of actions and interactions between point-light agents. European Journal of Neuroscience. https://doi.org/10.1111/EJN.15903

Haker, H., Schneebeli, M., & Stephan, K. E. (2016). Can Bayesian theories of autism spectrum disorder help improve clinical practice? Frontiers in Psychiatry, 7(JUN), 107. https://doi.org/10.3389/FPSYT.2016.00107/BIBTEX

Hallett, M. (2007). Transcranial Magnetic Stimulation: A Primer. Neuron, 55(2), 187–199. https://doi.org/10.1016/J.NEURON.2007.06.026

Harris, C. R., Millman, K. J., van der Walt, S. J., Gommers, R., Virtanen, P., Cournapeau, D., Wieser, E., Taylor, J., Berg, S., Smith, N. J., Kern, R., Picus, M., Hoyer, S., van Kerkwijk, M. H., Brett, M., Haldane, A., del Río, J. F., Wiebe, M., Peterson, P., … Oliphant, T. E. (2020). Array programming with NumPy. Nature 2020 585:7825, 585(7825), 357–362. https://doi.org/10.1038/s41586-020-2649-2

Hommel, B., Müsseler, J., Aschersleben, G., & Prinz, W. (2001). The Theory of Event Coding (TEC): a framework for perception and action planning. The Behavioral and Brain Sciences, 24(5), 849–878. https://doi.org/10.1017/S0140525X01000103

Huang, Y. Z., Edwards, M. J., Rounis, E., Bhatia, K. P., & Rothwell, J. C. (2005). Theta burst stimulation of the human motor cortex. Neuron, 45(2), 201–206. https://doi.org/10.1016/j.neuron.2004.12.033

Hunter, J. D. (2007). Matplotlib: A 2D Graphics Environment. Computing in Science & Engineering, 9(3), 90–95. https://doi.org/10.1109/MCSE.2007.55

Isik, L., Koldewyn, K., Beeler, D., & Kanwisher, N. (2017). Perceiving social interactions in the posterior superior temporal sulcus. Proceedings of the National Academy of Sciences of the United States of America, 114(43), E9145–E9152. https://doi.org/10.1073/PNAS.1714471114/SUPPL_FILE/PNAS.1714471114.SM04.AVI

JASP Team. (2022). JASP (Version 0.17)[Computer software]. https://jasp-stats.org/

Kilner, J. M., Vargas, C., Duval, S., Blakemore, S. J., & Sirigu, A. (2004). Motor activation prior to observation of a predicted movement. Nature Neuroscience, 7(12), 1299–1301. https://doi.org/10.1038/NN1355

Kleiner, M., Brainard, D., & Pelli, D. (2007). What’s new in Psychtoolbox-3?

Krol, M. A., Schutter, D. J. L. G., & Jellema, T. (2020). Sensorimotor cortex activation during anticipation of upcoming predictable but not unpredictable actions. Social Neuroscience, 15(2), 214–226. https://doi.org/10.1080/17470919.2019.1674688

Lapenta, O. M., Valasek, C. A., Vieira, S. M. G., & Boggio, P. S. (2022). Reading Point-Light Walkers and Amorphous—A TMS Study. Psychology and Neuroscience, 15(2), 198–209. https://doi.org/10.1037/PNE0000273

Lee, M. D., & Wagenmakers, E.-J. (2014). Bayesian Cognitive Modeling: A Practical Course. Cambridge University Press. https://doi.org/DOI:10.1017/CBO9781139087759

Manera, V., Becchio, C., Schouten, B., Bara, B. G., & Verfaillie, K. (2011a). Communicative interactions improve visual detection of biological motion. PLoS ONE, 6(1). https://doi.org/10.1371/journal.pone.0014594

Manera, V., del Giudice, M., Bara, B. G., Verfaillie, K., & Becchio, C. (2011b). The Second- Agent Effect: Communicative Gestures Increase the Likelihood of Perceiving a Second Agent. PLOS ONE, 6(7), e22650. https://doi.org/10.1371/JOURNAL.PONE.0022650

Manera, V., Schouten, B., Becchio, C., Bara, B. G., & Verfaillie, K. (2010). Inferring intentions from biological motion: A stimulus set of point-light communicative interactions. Behavior Research Methods, 42(1), 168–178. https://doi.org/10.3758/BRM.42.1.168

Maranesi, M., Livi, A., Fogassi, L., Rizzolatti, G., & Bonini, L. (2014). Mirror Neuron Activation Prior to Action Observation in a Predictable Context. Journal of Neuroscience, 34(45), 14827–14832. https://doi.org/10.1523/JNEUROSCI.2705-14.2014

Michael, J., Sandberg, K., Skewes, J., Wolf, T., Blicher, J., Overgaard, M., & Frith, C. D. (2014). Continuous Theta-Burst Stimulation Demonstrates a Causal Role of Premotor Homunculus in Action Understanding. Psychological Science, 25(4), 963–972. https://doi.org/10.1177/0956797613520608/ASSET/IMAGES/LARGE/10.1177_0956797 613520608-FIG2.JPEG

Miniussi, C., Ruzzoli, M., & Walsh, V. (2010). The mechanism of transcranial magnetic stimulation in cognition. Cortex, 46(1), 128–130. https://doi.org/10.1016/J.CORTEX.2009.03.004

Moss, F., Ward, L. M., & Sannita, W. G. (2004). Stochastic resonance and sensory information processing: a tutorial and review of application. Clinical Neurophysiology, 115(2), 267–281. https://doi.org/10.1016/J.CLINPH.2003.09.014

Neri, P., Morrone, M. C., & Burr, D. C. (1998). Seeing biological motion. Nature 1998 *395*:6705, *395*(6705), 894–896. https://doi.org/10.1038/27661

Ozdemir, R. A., Boucher, P., Fried, P. J., Momi, D., Jannati, A., Pascual-Leone, A., Santarnecchi, E., & Shafi, M. M. (2021). Reproducibility of cortical response modulation induced by intermittent and continuous theta-burst stimulation of the human motor cortex. Brain Stimulation, 14(4), 949–964. https://doi.org/10.1016/J.BRS.2021.05.013

Pavlova, M. A. (2012). Biological Motion Processing as a Hallmark of Social Cognition. Cerebral Cortex, 22(5), 981–995. https://doi.org/10.1093/CERCOR/BHR156

Pelli, D. G., & Vision, S. (1997). The VideoToolbox software for visual psychophysics: Transforming numbers into movies. Spatial Vision, 10, 437–442.

Pellicano, E., & Burr, D. (2012). When the world becomes “too real”: A Bayesian explanation of autistic perception. Trends in Cognitive Sciences, 16(10), 504–510. https://doi.org/10.1016/j.tics.2012.08.009

Ptak, R., Schnider, A., & Fellrath, J. (2017). The Dorsal Frontoparietal Network: A Core System for Emulated Action. Trends in Cognitive Sciences, 21(8), 589–599. https://doi.org/10.1016/J.TICS.2017.05.002

Rizzolatti, G., & Fadiga, L. (1998). Grasping objects and grasping action meanings: the dual role of monkey rostroventral premotor cortex (area F5). Novartis Foundation Symposium, 218(218), 81–103. https://doi.org/10.1002/9780470515563.CH6

Romero, M. C., Merken, L., Janssen, P., & Davare, M. (2022). Neural effects of continuous theta-burst stimulation in macaque parietal neurons. ELife, 11. https://doi.org/10.7554/ELIFE.65536

Rosenthal, R. (1994). Parametric Measures of Effect Size. In H. Cooper & L. V Hedges (Eds.), The Handbook of Research Synthesis (pp. 232–243). Russell Sage Foundation.

Rossi, S., Antal, A., Bestmann, S., Bikson, M., Brewer, C., Brockmöller, J., Carpenter, L. L., Cincotta, M., Chen, R., Daskalakis, J. D., Di Lazzaro, V., Fox, M. D., George, M. S., Gilbert, D., Kimiskidis, V. K., Koch, G., Ilmoniemi, R. J., Pascal Lefaucheur, J., Leocani, L., … Hallett, M. (2021). Safety and recommendations for TMS use in healthy subjects and patient populations, with updates on training, ethical and regulatory issues: Expert Guidelines. Clinical Neurophysiology : Official Journal of the International Federation of Clinical Neurophysiology, 132(1), 269–306. https://doi.org/10.1016/J.CLINPH.2020.10.003

Rossi, S., Hallett, M., Rossini, P. M., Pascual-Leone, A., Avanzini, G., Bestmann, S., Berardelli, A., Brewer, C., Canli, T., Cantello, R., Chen, R., Classen, J., Demitrack, M., Di Lazzaro, V., Epstein, C. M., George, M. S., Fregni, F., Ilmoniemi, R., Jalinous, R., … Ziemann, U. (2009). Safety, ethical considerations, and application guidelines for the use of transcranial magnetic stimulation in clinical practice and research. Clinical Neurophysiology : Official Journal of the International Federation of Clinical Neurophysiology, 120(12), 2008–2039. https://doi.org/10.1016/J.CLINPH.2009.08.016

Ruzzoli, M., Marzi, C. A., & Miniussi, C. (2010). The neural mechanisms of the effects of transcranial magnetic stimulation on perception. Journal of Neurophysiology, 103(6), 2982–2989. https://doi.org/10.1152/jn.01096.2009

Salge, J. H., Pollmann, S., & Reeder, R. R. (2020). Anomalous visual experience is linked to perceptual uncertainty and visual imagery vividness. Psychological Research 2020 *85*:5, *85*(5), 1848–1865. https://doi.org/10.1007/S00426-020-01364-7

Schilbach, L. (2016). Towards a second-person neuropsychiatry. Philosophical Transactions of the Royal Society B: Biological Sciences, 371(1686), 20150081. https://doi.org/10.1098/RSTB.2015.0081

Schilbach, L., Hoffstaedter, F., Müller, V., Cieslik, E. C., Goya-Maldonado, R., Trost, S., Sorg, C., Riedl, V., Jardri, R., Sommer, I., Kogler, L., Derntl, B., Gruber, O., & Eickhoff, S. B. (2016). Transdiagnostic commonalities and differences in resting state functional connectivity of the default mode network in schizophrenia and major depression. NeuroImage: Clinical, 10, 326–335. https://doi.org/10.1016/J.NICL.2015.11.021

Schubotz, R. I. (2007). Prediction of external events with our motor system: towards a new framework. Trends in Cognitive Sciences, 11(5), 211–218. https://doi.org/10.1016/J.TICS.2007.02.006

Schubotz, R. I., & Von Cramon, D. Y. (2002). A blueprint for target motion: fMRI reveals perceived sequential complexity to modulate premotor cortex. NeuroImage, 16(4), 920– 935. https://doi.org/10.1006/nimg.2002.1183

Schubotz, R. I., Wurm, M. F., Wittmann, M. K., & von Cramon, D. Y. (2014). Objects tell us what action we can expect: Dissociating brain areas for retrieval and exploitation of action knowledge during action observation in fMRI. Frontiers in Psychology, 5(JUN). https://doi.org/10.3389/fpsyg.2014.00636

Schwarzkopf, D. S., Silvanto, J., & Rees, G. (2011). Stochastic Resonance Effects Reveal the Neural Mechanisms of Transcranial Magnetic Stimulation. Journal of Neuroscience, 31(9), 3143–3147. https://doi.org/10.1523/JNEUROSCI.4863-10.2011

Siebner, H. R., Lang, N., Rizzo, V., Nitsche, M. A., Paulus, W., Lemon, R. N., & Rothwell, J. C. (2004). Preconditioning of low-frequency repetitive transcranial magnetic stimulation with transcranial direct current stimulation: evidence for homeostatic plasticity in the human motor cortex. The Journal of Neuroscience : The Official Journal of the Society for Neuroscience, 24(13), 3379–3385. https://doi.org/10.1523/JNEUROSCI.5316-03.2004

Spyder Developer Team. (2020). Spyder. https://www.spyder-ide.org/. Accessed 8 August 2021.

Stadler, W., Ott, D. V. M., Springer, A., Schubotz, R. I., Schütz-Bosbach, S., & Prinz, W. (2012). Repetitive TMS suggests a role of the human dorsal premotor cortex in action prediction. *Frontiers in Human Neuroscience*, FEBRUARY 2012. https://doi.org/10.3389/fnhum.2012.00020

Sterzer, P., Adams, R. A., Fletcher, P., Frith, C., Lawrie, S. M., Muckli, L., Petrovic, P., Uhlhaas, P., Voss, M., & Corlett, P. R. (2018). The Predictive Coding Account of Psychosis. Biological Psychiatry, 84(9), 634–643. https://doi.org/10.1016/J.BIOPSYCH.2018.05.015

Tarasi, L., Magosso, E., Ricci, G., Ursino, M., & Romei, V. (2021). The directionality of fronto- posterior brain connectivity is associated with the degree of individual autistic traits. Brain Sciences, 11(11). https://doi.org/10.3390/brainsci11111443

Tarasi, L., Trajkovic, J., Diciotti, S., di Pellegrino, G., Ferri, F., Ursino, M., & Romei, V. (2022). Predictive waves in the autism-schizophrenia continuum: A novel biobehavioral model. In Neuroscience and Biobehavioral Reviews (Vol. 132, pp. 1–22). Elsevier Ltd. https://doi.org/10.1016/j.neubiorev.2021.11.006

Umiltà, M. A., Kohler, E., Gallese, V., Fogassi, L., Fadiga, L., Keysers, C., & Rizzolatti, G. (2001). I know what you are doing: A neurophysiological study. Neuron, 31(1), 155–165. https://doi.org/10.1016/S0896-6273(01)00337-3

Urgesi, C., Maieron, M., Avenanti, A., Tidoni, E., Fabbro, F., & Aglioti, S. M. (2010). Simulating the Future of Actions in the Human Corticospinal System. Cerebral Cortex, 20(11), 2511–2521. https://doi.org/10.1093/CERCOR/BHP292

van de Cruys, S., Evers, K., van der Hallen, R., van Eylen, L., Boets, B., de-Wit, L., & Wagemans, J. (2014). Precise minds in uncertain worlds: Predictive coding in autism. Psychological Review, 121(4), 649–675. https://doi.org/10.1037/A0037665

van Kemenade, B. M., Muggleton, N., Walsh, V., & Saygin, A. P. (2012). Effects of TMS over Premotor and Superior Temporal Cortices on Biological Motion Perception. Journal of Cognitive Neuroscience, 24(4), 896–904. https://doi.org/10.1162/JOCN_A_00194

Van Rossum G, & Drake, F. L. (2009). Python 3 reference manual. CreateSpace, Scotts Valley.

Vanrie, J., Dekeyser, M., & Verfaillie, K. (2016). Bistability and Biasing Effects in the Perception of Ambiguous Point-Light Walkers. Http://Dx.Doi.Org/10.1068/P5004, 33(5), 547–560. https://doi.org/10.1068/P5004

Varnava, A., Stokes, M. G., & Chambers, C. D. (2011). Reliability of the “observation of movement” method for determining motor threshold using transcranial magnetic stimulation. Journal of Neuroscience Methods, 201(2), 327–332. https://doi.org/10.1016/J.JNEUMETH.2011.08.016

Varrier, R. S., & Finn, E. S. (2022). Seeing Social: A Neural Signature for Conscious Perception of Social Interactions. Journal of Neuroscience, 42(49), 9211–9226. https://doi.org/10.1523/JNEUROSCI.0859-22.2022

Veale, J. F. (2013). Edinburgh Handedness Inventory – Short Form: A revised version based on confirmatory factor analysis. Http://Dx.Doi.Org/10.1080/1357650X.2013.783045, 19(2), 164–177. https://doi.org/10.1080/1357650X.2013.783045

von der Lühe, T., Manera, V., Barisic, I., Becchio, C., Vogeley, K., & Schilbach, L. (2016). Interpersonal predictive coding, not action perception, is impaired in autism. Philosophical Transactions of the Royal Society B: Biological Sciences, 371(1693). https://doi.org/10.1098/RSTB.2015.0373

Walsh, V., & Cowey, A. (2000). Transcranial magnetic stimulation and cognitive neuroscience. Nature Reviews. Neuroscience, 1(1), 73–80. https://doi.org/10.1038/35036239

Wheaton, K. J., Thompson, J. C., Syngeniotis, A., Abbott, D. F., & Puce, A. (2004). Viewing the motion of human body parts activates different regions of premotor, temporal, and parietal cortex. NeuroImage, 22(1), 277–288. https://doi.org/10.1016/J.NEUROIMAGE.2003.12.043

Zillekens, I. C., Brandi, M. L., Lahnakoski, J. M., Koul, A., Manera, V., Becchio, C., & Schilbach, L. (2019). Increased functional coupling of the left amygdala and medial prefrontal cortex during the perception of communicative point-light stimuli. Social Cognitive and Affective Neuroscience, 14(1), 97–107. https://doi.org/10.1093/scan/nsy105

